# MPI-Guided Photothermal Therapy of Prostate Cancer using Stem Cell Delivery of Magnetotheranostic Nanoflowers

**DOI:** 10.1101/2025.04.27.650870

**Authors:** Behnaz Ghaemi, Ana Rosu, Shreyas Kuddannaya, Gautier Laurent, Rana Bazzi, Stéphane Roux, Ali Shakeri-Zadeh, Jeff W. M. Bulte

## Abstract

Intratumoral nanoparticle (NP) injection is a commonly used for local hyperthermia. A limitation of this delivery route is the limited NP dispersion and retainment within the tumor, with undesired leakage leading to off-target toxicity. By exploiting their inherent tumor tropism, we have used human mesenchymal stem cells (hMSCs) as delivery vehicles for magnetic theranostic gold-iron oxide nanoflowers (GIONF) to improve their overall intratumoral distribution and retention. GIONF-loaded hMSCs exhibited excellent heating properties for laser photothermal therapy and high tracer performance for visualization and quantification by magnetic particle imaging (MPI). In contrast to naked GIONF, GIONF-hMSCs remained within the prostate tumors one week post-injection, with MPI-guided PTT completely ablating tumors without recurrence.

---

Imaging-guided photothermal therapy (PTT) employs photosensitizing nanoparticles (NPs) as imaging agents that convert laser into heat capable of killing cancer cells^1^. Gold nanoparticles have been most commonly used due to their favorable properties, which includes their small size for tissue penetration, overall biocompatibility, and efficient laser-heat conversion in the near-infrared (NIR) region^2, 3^. Systemic injection of NPs is often inadequate as only a small % of the injected dose ends up in the tumor, with most particles localizing in cells belonging to the reticulo-endothelial system^4–6^. Direct intratumoral (i.t.) injection of NPs is far more effective but could also lead to systemic circulation and off-target accumulation if backflow leakage or poor tumor retention occurs. A promising approach to enhance the distribution and retention of NPs at the tumor site is to employ tumor-tropic human mesenchymal stem cells (MSCs) as a smart biovehicle^7–10^. The potential of tumor homing of hMSCs is ascribed to the presence of surface-associated chemokine receptors, such as CXCR4, and their binding to chemokines secreted by the tumor^8, 11^. However, be it in naked form or loaded into cells, uncertainties about tumor retention and overall body distribution of NPs will remain, hampering interpretations related to therapeutic outcome. Hence, it is desirable to develop non-invasive imaging techniques that can report on the pharmacokinetics of NPs for imaging-guided PTT^12^.

We have employed a hybrid magnetotheranostic nanoflower that can be used for simultaneous PTT and magnetic particle imaging (MPI) by virtue of their gold and iron oxide content, respectively, termed gold iron oxide nanoflowers (GIONF). MPI is a quantitative imaging modality^13^, that, when combined with CT as anatomical imaging modality, can report on the local and whole-body distribution of administered naked GIONF and GIONF loaded in hMSCs. As a proof of principle, we chose prostate cancer as tumor model since it is readily accessible via either the urethra or rectum to perform biopsies and endoscopic procedures including laser irradiation in patients^14^. We hypothesized that GIONF-labeled hMSCs could be attractive candidates for tumor PPT as the cells distribute and retain GIONF throughout the entire tumor, which can not only be visualized but also quantified with MPI^13^. Since GIONF-labeled hMSCs are destroyed inside the tumor upon completion of PTT, any potential risks of hMSC-mediated tumor neoangiogenesis and/or other stromal effects promoting tumor growth^15^ can be avoided. After comparing the i.v. and i.t routes of injection using MPI/CT, the imaging data were used to guide on-demand laser triggering at the time point of maximum retention and distribution of GIONF-hMSCs within the tumor.

## GIONF labeling of hMSCs

GIONF is composed of an iron oxide nanoflower core, 30 nm in size, grafted with ultrasmall gold NPs measuring 2-3 nm in diameter. GIONF has continuous crystalline orientations, which enhances the effective particle magnetic moment and performance. Prussian Blue staining confirmed intracellular uptake of GIONFs in hMSCs near the perinuclear region, that was higher when poly-L-lysine (PLL) was added as polymeric cation (**Fig. 1A, S1**). An LDH assay demonstrated that GIONF-PLL labeling was not toxic to hMSCs at up to concentrations of at least 100 μg Fe/mL (**Fig. 1B**). A Ferrozine assay quantified an average of 40 pg Fe per hMSC (**Fig. 1C**). The MPI signal intensity showed a linear correlation (R^2^=0.998) with the number of GIONF-hMSCs (**Fig. 1D**), confirming its applicability for further *in vivo* quantification^5, 16^. MPI could detect as few as 5×10^3^ GIONF-hMSCs, indicating exquisite sensitivity for further *in vivo* imaging.

**Figure 1:**
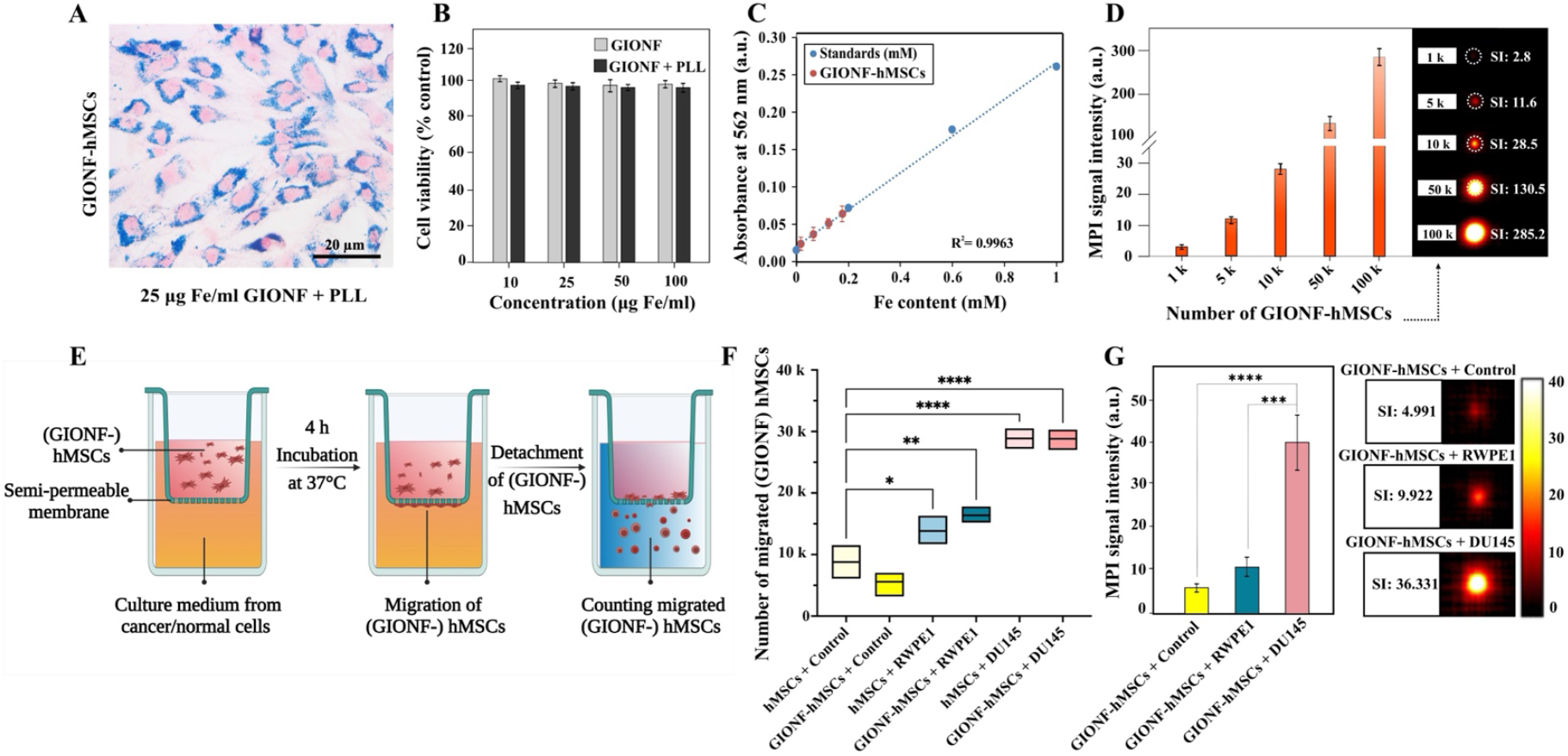
Intracellular labeling and migration properties of GIONF-labeled hMSCs. (**A**) Prussian Blue staining of hMSCs incubated for 24h with 25 μg Fe/mL of GIONF and 3.1 μg/ml of PLL shows NP accumulation within the cytoplasm. (**B**) hMSC viability after 24h incubation with 10-100 μg Fe/mL GIONF with and without 3.1 μg/ml PLL. (**C**) Calibration curve for calculating the iron content of GIONF-hMSCs. (**D**) MPI signal intensity and images of GIONF-hMSCs suspended in agarose gel. (**E**) Schematic of Boyden chamber assay for assessing hMSC tumor tropism. GIONF-hMSCs and unlabeled hMSCs were placed in the upper well, and the lower well was filled with prostate cancer cell line DU145 or normal prostate epithelial cell line RWPE1-conditioned media, or DMEM supplemented with FBS (control). After 4h of incubation, migrated hMSCs were detached from the well bottom and counted using confocal microscopy. (**F**) Average number of migrated hMSCs per well quantified by Luna digital cell counter (n=5 independent experiments). (**G**) MPI signal intensity and images of migrated hMSCs in 0.5% w/v agarose gel (n=6 independent experiments).

GIONF-hMSCs must retain their tumor tropism post-GIONF internalization to function as effective delivery agents. Iron oxide labeling can transiently reduce hMSC migration *in vitro* and *in vivo*, but this effect may diminish following multiple cell divisions and/or NP exocytosis^17, 18^. Unlabeled and GIONF-hMSCs were exposed for 4h to either prostate cancer cell line DU145- or normal prostate epithelial cell line RWPE1-conditioned media, or DMEM with FBS as control, using a Boyden chamber (**Fig. 1E**). Unlabeled hMSCs and GIONF-hMSCs had a significantly higher (six-fold, p<0.0001) migration towards DU145 and a three-fold higher (p< 0.01) migration towards RWPE1 cells compared to DMEM/FBS control (**Fig. 1F**). Bright field microscopy of cells that adhered to the bottom well corroborated these findings (**Fig. S2**). MPI furthermore confirmed the relative tumor tropism of GIONF-labeled hMSCs cells which quantitatively corresponded to the Boyden chamber assay cell counts, with an approximately 5-6- and 2-4-fold increase in signal for DU145- and RWPE1-conditioned media, respectively, compared to control (**Fig. 1G**). Importantly, GIONF labeling did not impair tumor tropism, as cell migrated cell numbers remained comparable between GIONF-hMSCs and unlabeled hMSCs (**Fig. 1F**).

### *In vivo* MPI of tumors injected with GIONF-hMSCs or naked GIONFs

MPI, with its high sensitivity and linear iron quantification, enables precise monitoring of GIONF distribution and retention. DU145 tumor-bearing mice received intravenous (i.v.) or intratumoral (i.t.) injections of 2.5×10^5^ GIONF-hMSCs (containing 40 pg Fe per cell equivalent to 10 μg total Fe) or 10 μg Fe naked GIONFs, followed by *in vivo* MPI at 24-hour intervals for one week (**Fig. 2, S3**). I.v. injection of naked GIONFs resulted in liver accumulation with no detectable tumor signal. The liver MPI signal gradually declined but remained detectable for seven days (**Fig. 2A**). I.v.-injected GIONF-hMSCs initially accumulated in the lungs before redistributing to the liver within 24h, with liver signal increasing at 24h while a reduced lung signals persisted (**Fig 2B and S3**). *Ex vivo* imaging confirmed these findings, showing sustained localization in both the lung and liver, with consistently stronger liver signals throughout the monitoring period without detectable MPI signal in the tumor (**Fig. S4**).

**Figure 2:**
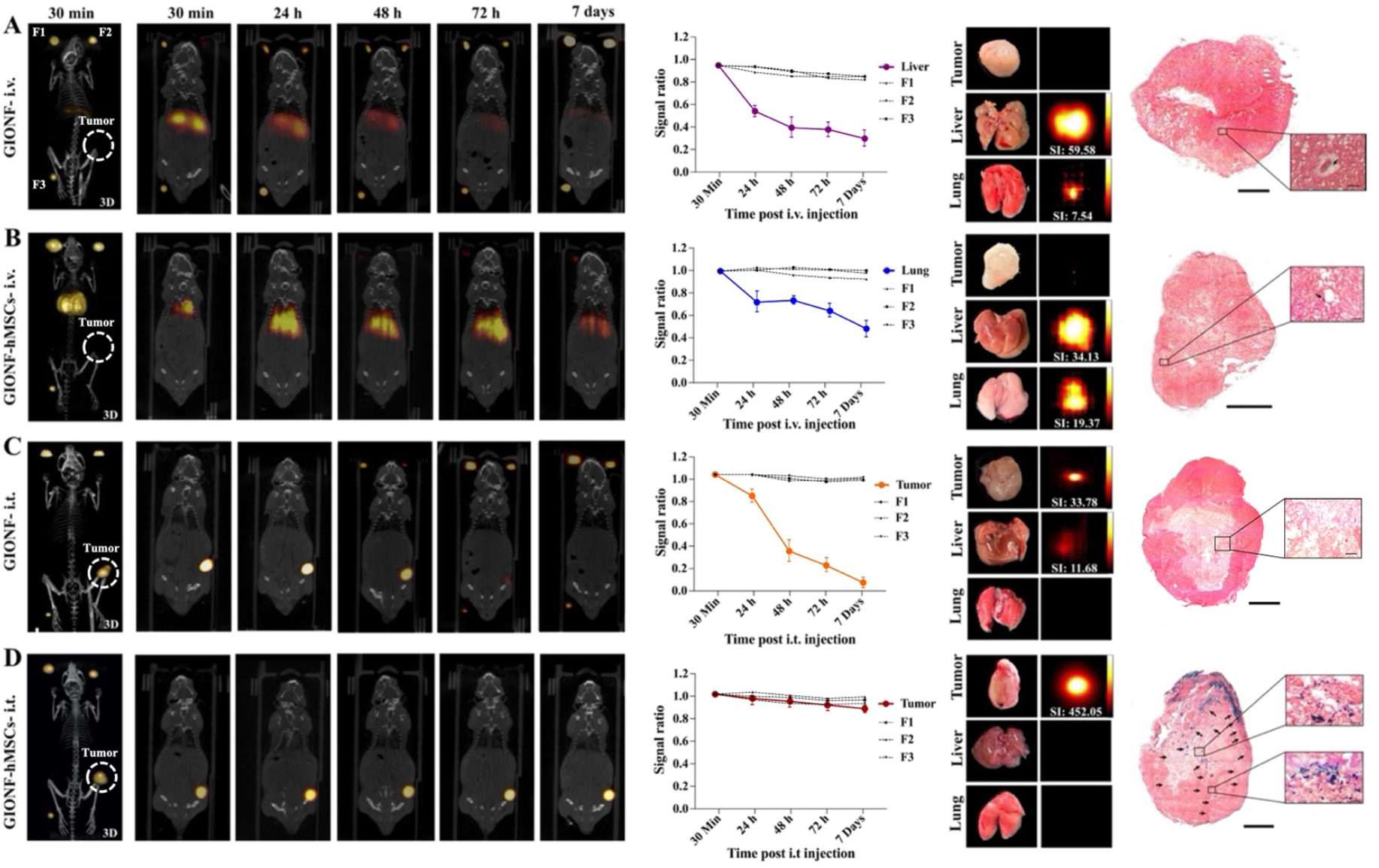
MPI/CT of naked GIONF and GIONF-hMSCs injected i.v. or i.t. in DU145 tumor-bearing mice. Shown are *in vivo* 3D and 2D MPI/CT images taken at several time points post (**A**,**B**) i.v. or (**C**,**D**) i.t. injection of (**A**,**C**) naked GIONF or (**B**,**D**) GIONF-hMSCs. Three fiducials (F1, F2, F3) containing 20, 10, and 5% of total injected GIONF-hMSCs were used as MPI signal calibration reference standards. The MPI signal ratio of the prostate tumor and fiducials was calculated by dividing the MPI signal intensity at each time point by the initial MPI signal intensity at 30 min post-injection, averaged for n=4 animals. 3D *ex vivo* MPI images of the tumor, liver and lung 7 days post-injection with corresponding PB-stained histological sections are shown in the right two columns. Scale bar=1000 μm.

The off-target localization of GIONFs following i.v. injection of both naked GIONFs and GIONF-hMSCs necessitates i.t. administration as an alternative. I.t.-injected naked GIONFs exhibited a ten-fold reduction in MPI signal intensity within 72h and were undetectable at the tumor site by day 7 (**Fig. 2C**). NP clearance rates are influenced by particle size, surface charge, and tumor composition^5, 19, 20^.

2D and 3D MPI showed that GIONF-hMSCs retained most of their initial MPI signal intensity at the tumor site when injected i.t, with only a 10% reduction seven days post injection with no detectable off-target accumulation (**Fig. 2D)**. To the best of our knowledge, this is the first study demonstrating the sustained retention of i.t.-injected NP-labeled hMSCs within the tumor beyond three days post-injection^21–23^.

Efficient NP distribution within the tumor is critical for optimal therapeutic outcomes especially for focal therapies such as PTT, where efficacy depends on NP localization. *Ex vivo* MPI performed seven days post-i.v. and i.t. injection of GIONF-hMSCs and naked GIONFs confirmed the *in vivo* MPI findings for off-target accumulation (**Fig. 2**). To analyze the microscopic NP distribution in the tumors, they were excised seven days post-injection and stained with PB for iron. Minimal GIONF accumulation was detected in tumors following i.v. injection, primarily near blood vessels. I.t. injection of GIONF-hMSCs led to widespread peripheral tumor distribution, whereas no iron deposition was observed in tumors injected with free GIONF. The tumor-tropism of hMSCs likely accounts for this extended retention and distribution, driven by cytokines, chemokines, and factors such as HIF. IL-6, IL-1β, IFN-γ, and TNF-α facilitate hMSC adhesion and tumor infiltration, while SDF-1/CXCR4 signaling promotes retention at the tumor periphery^8^, an area associated with aggressive tumor growth and metastasis. In contrast, the necrotic tumor core lacks key cytokines and receptors for hMSC homing. As a result, GIONF-hMSCs function as ‘smart’ biovehicles, delivering NPs to the tumor’s most aggressive regions—an outcome unachievable with naked NPs, which typically remain at the injection site or within the extracellular matrix before clearance^24 25, 26^.

### *In vitro* PTT of prostate tumor spheres incubated with naked GIONF or GIONF-hMSCs

The photothermal efficacy of GIONF-hMSCs and naked GIONFs was first evaluated *in vitro* using DU145 3D tumor spheres (**Fig. 3A**). Compared to 2D monolayer cultures, tumor spheres more accurately model dynamic tumor processes, including growth kinetics, differential gene expression, nutrient and oxygen gradients, and catabolite accumulation^27, 28^. Similar to *in vivo* tumors, they exhibit exponential proliferation followed by a growth plateau, where increasing nonproliferative and necrotic regions emerge due to diffusion limitations. In spheres >500 μm in diameter, an outer proliferative layer, an inner quiescent zone, and a necrotic core develop (**Fig. S5**). Before incubation with tumor spheres, GIONF-hMSCs and naked GIONFs were irradiated to assess and compare their inherent photothermal potential^29^. Given that GIONFs undergo physicochemical modifications and aggregation changes post-cellular uptake, comparing the heating capacity of naked GIONFs to intracellular GIONFs is crucial^30^. GIONF-hMSCs containing increasing GIONF amounts (0.65–10 μg Fe) were irradiated, with the highest amount reaching 60°C (**Fig. 3B**). Under the same conditions, naked GIONFs reached 42.7°C for 10 μg Fe (**Fig. S6**). GIONF-hMSCs exhibited enhanced heating efficiency compared to naked GIONFs for equivalent GIONF amounts. A similar effect was reported by Nicolas-Boluda et al. in EGI-1 cholangiocarcinoma cells, suggesting that intracellular confinement enhances GIONF heating capacity. This may be due to the nanoflower shape of GIONF and its clustering within endo-lysosomal compartments, enhancing radiation-to-heat conversion for efficient PTT by promoting plasmon coupling and shifting the optical absorption toward the NIR region, increasing energy absorption under NIR laser irradiation^29^. Compared to spherical SPIOs or gold NPs, GIONF exhibits two-fold higher magnetization and ten-fold greater heating capacity^29, 31^. In comparison, 1 mg/ml of naked GIONFs exposed to an 808 nm laser at 2.2 W/cm^2^ for 5 minutes reached a maximum temperature increase of 44 °C^29^, 0.5 mM magnetic gold nanowreaths exposed to NIR laser at 0.75 W/cm^2^ for 120 seconds reached 55 °C^32^, and 4 nM of gold-coated iron oxide nanoroses irradiated at 1.3 W/cm^2^ for 3 minutes achieved a maximum temperature of 50 °C^33^.

**Figure 3:**
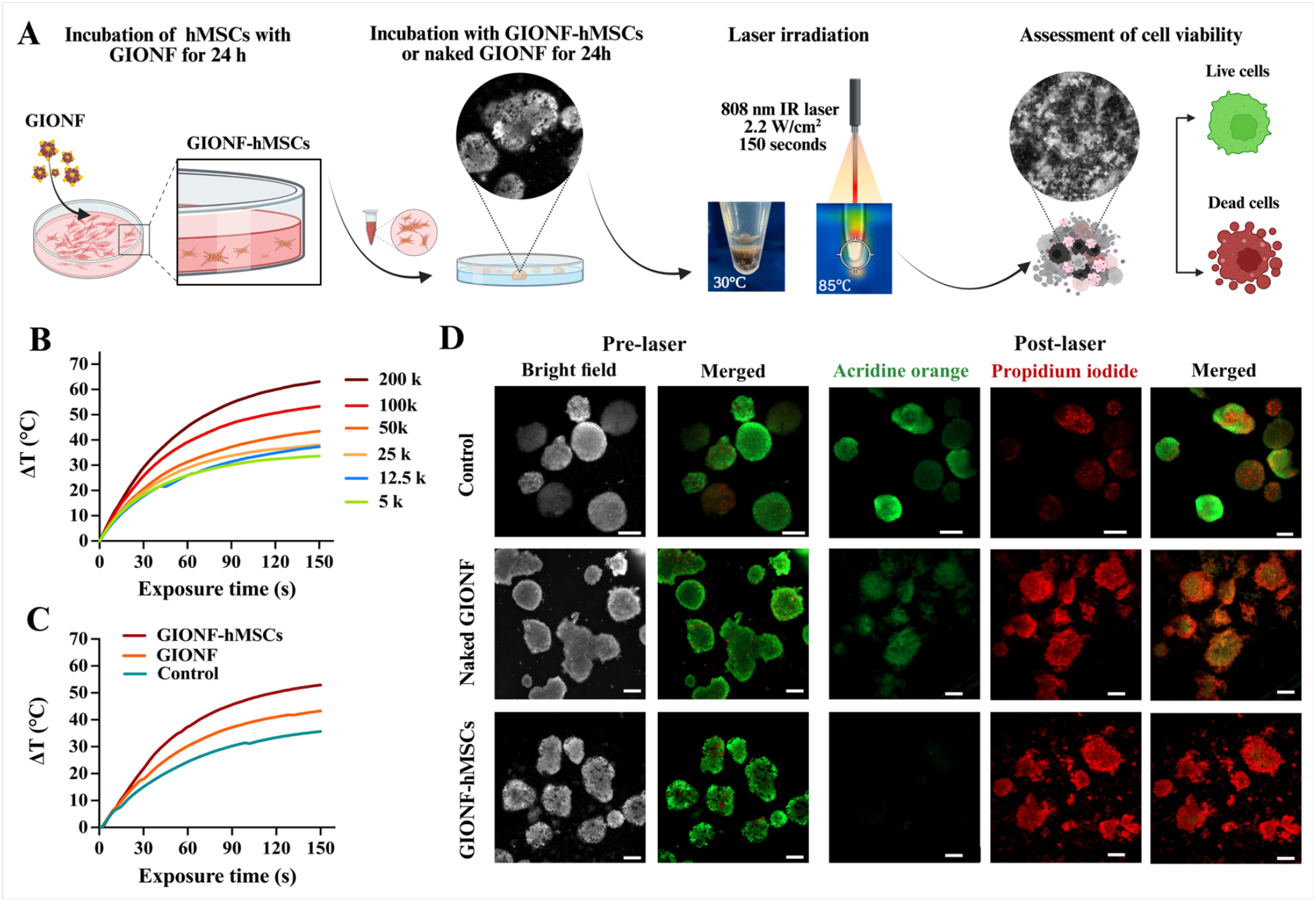
*In vitro* PTT studies of DU145 tumor spheres incubated with naked GIONF or GIONF-hMSCs. (**A**) Schematic illustration of experimental setup. (**B**) Thermometry of GIONF-hMSCs exposed to laser irradiation for 150 seconds. (**C**) Thermometry of DU145 spheres incubated with naked GIONFs of GIONF-hMSCs. (**D**) Acridine orange (AO) and propidium iodide (PI) staining of DU145 spheres incubated with naked GIONF or GIONF-hMSCs before and after laser irradiation. Control represents no incubation. AO and PI represent live and dead cells, respectively. Bar=200 μm.

DU145 tumor spheres were then incubated for 24h with naked GIONF or GIONF-hMSCs, followed by 808 nm NIR laser irradiation at 2.2 W/cm^2^ for 150 s with real-time temperature monitoring (**Fig. 3C**). Spheres treated with GIONF-hMSCs reached ~85°C, 10°C higher than those with naked GIONFs, while controls increased to 65°C. AO/PI staining confirmed complete tumor ablation with GIONF-hMSCs (**Fig. 3D**), whereas naked GIONFs induced only peripheral ablation, likely due to limited NP penetration and/or enhanced heating by GIONF-hMSCs^34^.

### *In vivo* MPI-guided PTT of prostate tumors injected i.t. with naked GIONF or GIONF-hMSCs

To evaluate the photothermal efficacy of GIONF-hMSCs, Rag2 mice bearing luciferase-positive DU145 (Nanoluc-DU145) tumors were divided into four experimental groups: (1) tumors injected i.t. with naked GIONFs and irradiated 30 minutes post-injection, (2) tumors injected i.t. with naked GIONFs and irradiated 48h post-injection, (3) tumors injected i.t. with GIONF-hMSCs and irradiated 48h post-injection, and (4) control tumors injected i.t. with PBS and irradiated to assess the effects of laser exposure alone. All groups underwent 808 nm laser irradiation at 2.2 W/cm^2^ for 7 min with real-time thermal imaging and thermometry.

The 48-hour time point for tumor imaging and irradiation was selected based on findings by Mooney et al., who demonstrated that stem cell-mediated delivery of nanoparticles achieves a significantly broader and more uniform intratumoral distribution within this timeframe compared tonaked nanoparticles, which remain largely confined to the injection site^35^. Naked GIONFs injected into tumors were irradiated 30 minutes post-injection, corresponding to their peak concentration within the tumor; in a separate group, naked GIONFs were reinjected at 48h to serve as controls for nanoparticle retention and distribution in comparison to stem cell– mediated delivery.

Following irradiation, tumors reached different peak temperatures: 42.0 °C in control tumors, 51.1 °C in tumors injected with naked GIONFs and irradiated 30 minutes post-injection, 48.4 °C in tumors injected with naked GIONFs and irradiated 48h post-injection, and the highest temperature of 57.9 °C in tumors injected with GIONF-hMSCs and irradiated 48h post-injection (**Fig. 4A)**. The greater temperature rise observed with GIONF-hMSCs is attributed to their widespread i.t. tumor distribution, which facilitates more efficient heat transfer. In contrast, naked GIONFs remained localized at the injection site, leading to slower, localized heating with a continuous temperature increase over seven minutes of irradiation (**Fig. 4B**). MPI confirmed that naked GIONFs began clearing from the tumor by 48h post-injection, explaining the lower temperature increase at this time point. Thermal thresholds for tumor ablation further distinguished treatment efficacy. Control tumors failed to reach the necrosis threshold (42°C for 10 minutes), while tumors injected with naked GIONFs reached 48–51.1°C, leading to partial tumor ablation via microvascular thrombosis and ischemia^1^. In contrast, GIONF-hMSCs achieved temperatures above 57.9°C, resulting in near-instantaneous tumor ablation due to protein denaturation and plasma membrane disruption^1^. These findings indicate that naked GIONFs alone are insufficient for complete tumor destruction, whereas GIONF-hMSCs efficiently distribute nanoparticles throughout the tumor, significantly enhancing photothermal efficiency and achieving complete tumor ablation^36^.

**Figure 4:**
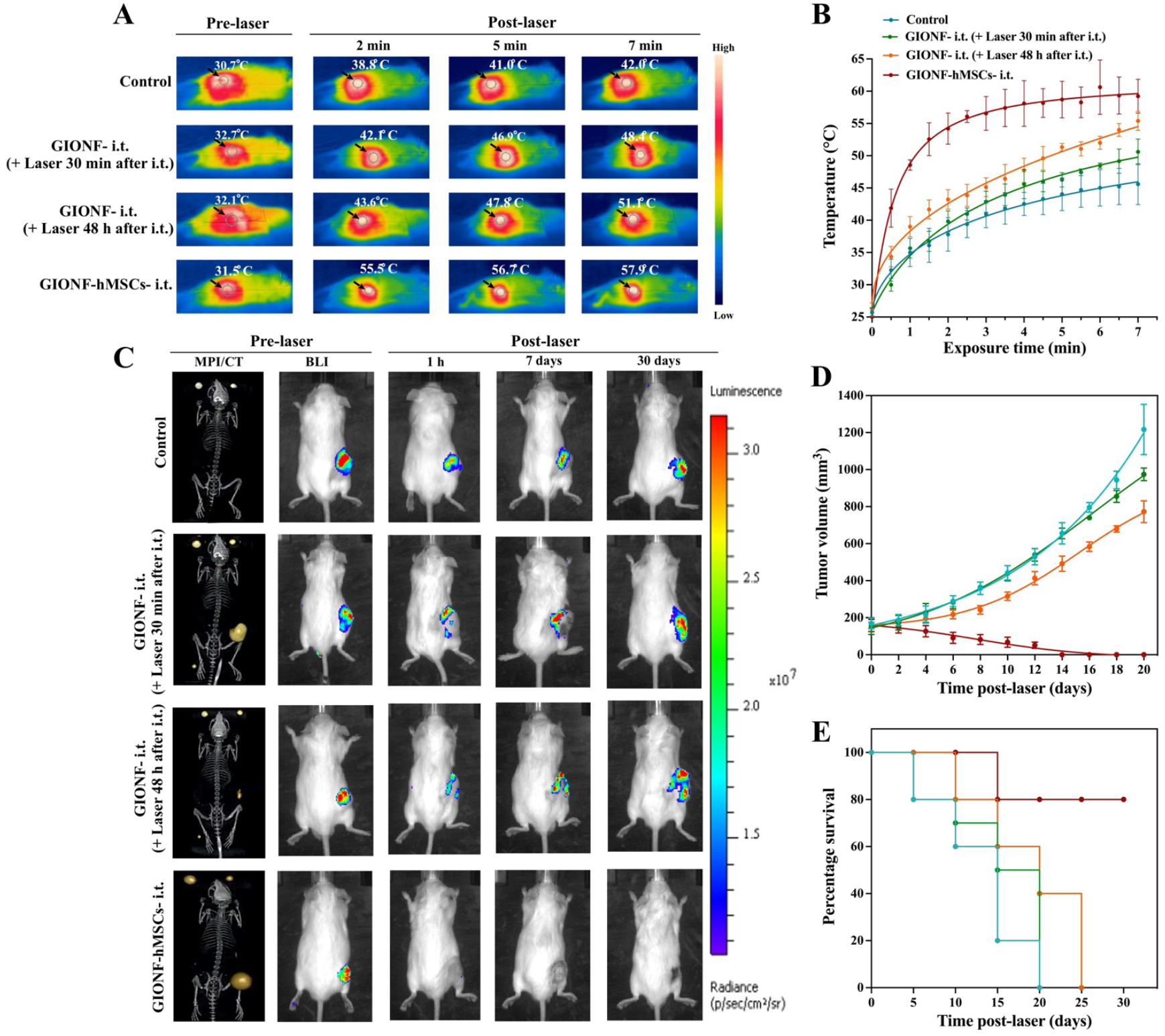
*In vivo* PTT and BLI of DU145 bearing mice injected i.t. with naked GIONF or GIONF-hMSCs. (**A**) Thermal images and (**B**) temperature quantification of DU145 tumor xenografts injected with naked GIONF or GIONF-hMSCs pre- and post-laser irradiation. Irradiation was performed either 30 min or 48 h post injection. Control = no injection (laser irradiation only). (**C**) Serial BLI of DU145 xenografts injected with naked GIONF or GIONF-hMSCs pre- and post-laser irradiation. Irradiation was performed either 30 min or 48 h post injection. Control = no injection (laser irradiation only). (**D**) Tumor volume measurements and (**E**) percentage animal survival for control mice and mice injected i.t. with naked GIONF of GIONF-hMSCs. Data represent mean values±SD (n=3 animals per group).

We used MPI and bioluminescence imaging (BLI) to assess the i.t.-injected GIONF content pre-PTT and tumor viability post-PTT, respectively (**Fig. 4C**). MPI signals confirmed equivalent GIONF quantities in tumors injected i.t. with naked GIONFs (imaged at 30 minutes post-injection) and those injected i.t. with GIONF-hMSCs. However, tumors injected with naked GIONFs exhibited a significantly reduced MPI signal by 48h post-injection, indicating nanoparticle clearance. BLI showed that control tumors remained viable for up to 30 days post-PTT, confirming that transient heating to 42°C was insufficient for tumor ablation. Naked GIONF-treated tumors exhibited greater cell death when irradiated 30 minutes post-injection compared to 48h post-injection, though some peripheral tumor cells survived. By seven days post-PTT, both groups showed tumor recurrence, with complete regrowth at day 30. In contrast, GIONF-hMSC-treated tumors exhibited complete ablation within 1 hour post-PTT, with no recurrence up to 30 days. Burned tumor fragments detached between days 12 and 16 post-PTT, further confirming effective ablation (**Fig. 4D**). Tumors treated with naked GIONFs reached 1000 mm^3^, representing a 5-fold increase in initial tumor volume. Survival rates correlated with BLI and tumor volume trends, with mice injected with GIONF-hMSCs achieving 80% survival at 90 days post-PTT (**Fig. 4E**).

The efficacy of hMSCs as carriers for enhancing PTT has also been demonstrated for various other nanoparticles. Breast tumor xenografts injected i.v. with LDGI-labeled hMSCs (lipid, doxorubicin, gold nanorod, and iron oxide nanoclusters) reached a final temperature of 59.9°C, significantly higher than the 48.6°C observed for naked LDGI^22^. This difference may be attributed to tumor-tropic hMSCs delivering more LDGI to the tumor compared to passive accumulation of naked LDGI. Similarly, fibrosarcoma xenografts treated with pH-sensitive gold nanoparticle (PSAuNP)-labeled hMSCs exhibited a 6.3°C higher temperature increase than naked PSAuNPs^37^, comparable to the temperature difference observed between GIONF-hMSCs and naked GIONFs in our study.

### Biodistribution studies and histopathology

H&E staining and TUNEL assays were performed on excised tumors to assess the mechanism of cell death post-PTT (**Fig. 5A**). One-hour post-PTT, control tumors exhibited minimal apoptotic activity, primarily at the periphery, with most cells remaining viable. Tumors irradiated 30 minutes post-i.t. injection of naked GIONFs displayed localized apoptosis at the injection site, while i.t. injection of GIONF-hMSCs resulted in the most homogeneous apoptotic activity. By seven days post-PTT, tumors injected with naked GIONF maintained localized apoptosis, with viable cells persisting in the core and most of the periphery. In contrast, GIONF-hMSC-treated tumors exhibited widespread apoptosis, corresponding to complete tumor ablation. By 30 days post-PTT, scar tissue formation was observed in the GIONF-hMSC group, and by day 90, complete healing occurred without any signs of recurrence (**Fig. 5B)**.

**Figure 5:**
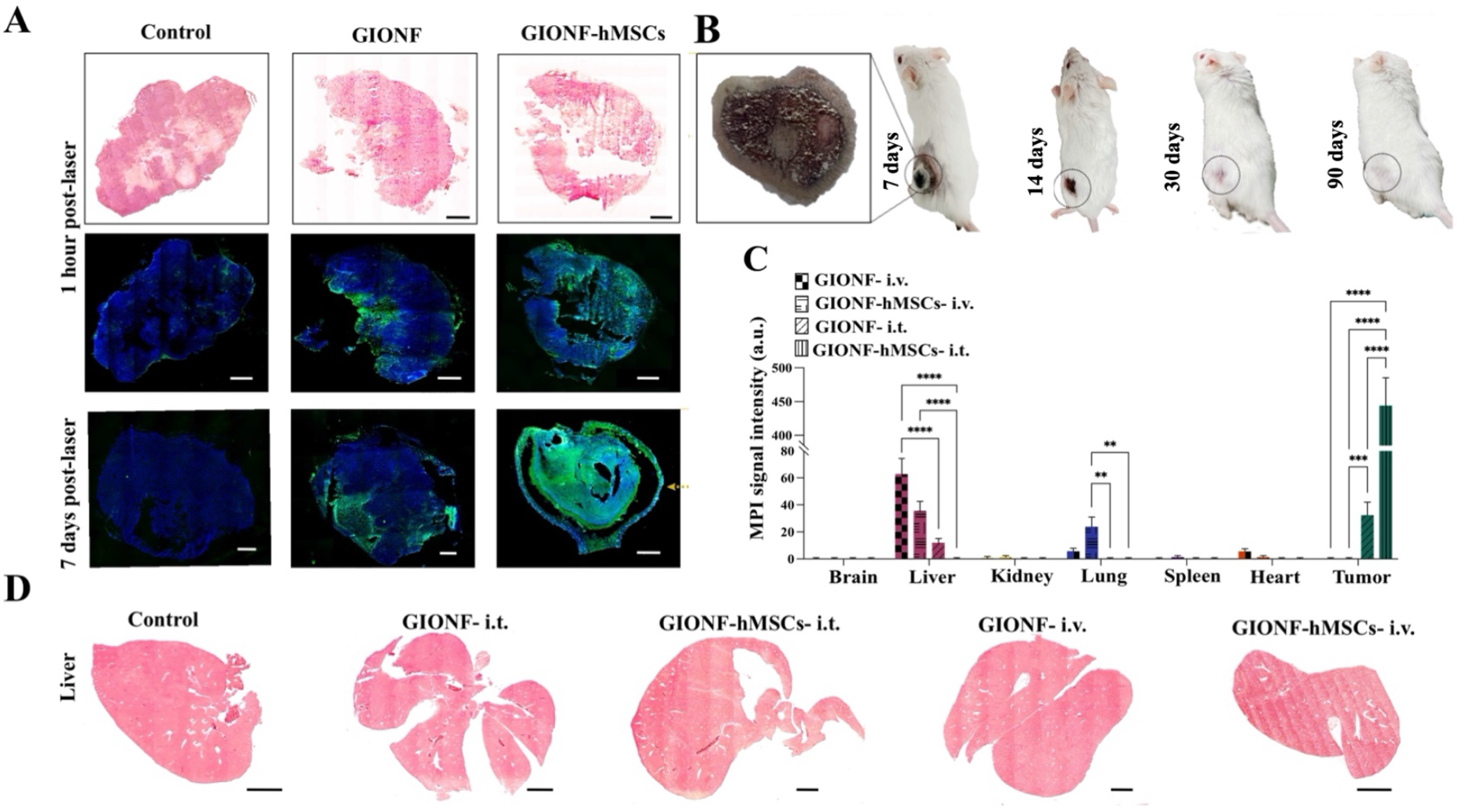
Post-mortem histology of DU145 prostate tumors injected i.t. or i.v. with naked GIONF, GIONF-hMSCs or no injection (control), subjected to PTT. (**A**) H&E staining and TUNEL assay 1h and 7 days post-PTT. Blue (DAPI) marks cell nuclei; green represents TUNEL-positive apoptotic cells showing DNA fragmentation (**B**) Photograph of ablated tumor section injected with GIONF-hMSCs 7 days post-PTT and disappearance of the tumor in a representative mouse over 90 days. (**C**) Quantitative *ex vivo* MPI signals defining biodistribution of GIONF in vital organs 7 days post i.v. and i.t injection of naked GIONF and GIONF-hMSCs (**D**) H&E staining of the mice livers, 7 days post i.v. and i.t injection of naked GIONF and GIONF-hMSCs compared to control. Scale bar=1 mm.

Molecular analysis of p32 and caspase-3 in prior studies indicated nearly 100% apoptosis throughout the tumor following PTT with MSC-PSAuNPs, compared to 40% apoptosis with naked PSAuNPs^38^. To assess the biodistribution and potential toxicity of GIONFs administered via different routes, *ex vivo* MPI quantification was performed on the major organs seven days post-injection. Following *i*.*v*. injection, naked GIONF exhibited strong MPI signals in the liver, with minimal signals in the lung and heart and no detectable tumor signal. In contrast, i.v.-injected GIONF-hMSCs showed significant MPI signals in both the liver and lung.

For i.t. injection, naked GIONFs were detected in both the tumor and liver, suggesting translocation over time, whereas GIONF-hMSCs exhibited strong tumor retention with no detectable off-target accumulation, confirming their tumor-tropic properties (**Fig. 5C**). Despite prolonged liver retention of NPs, which may raise concerns regarding chronic toxicity, H&E staining revealed no significant liver damage in any of the experimental groups (**Fig. 5D**). These findings support the biocompatibility and safety profile of GIONF and GIONF-hMSCs, reinforcing their potential for clinical applications. Despite prolonged hepatic retention, iron oxide NPs exhibit no toxicity unless administered i.v. at very high doses (≥10 mg/kg)^39 39^, while gold NPs can alter gene expression related to detoxification and metabolism^40^. hMSC-based NP delivery may mitigate these effects by retaining NPs within tumors, reducing systemic exposure, and slowing their release.

## Conclusion

We demonstrated the effectiveness of GIONF-labeled hMSCs as a cellular nanotheranostic MPI/PTT agent for prostate cancer with improved retention and intratumoral distribution compared to naked GIONF. A single-shot i.t. injection of GIONF-hMSCs was able to completely ablate tumors with no recurrence after 90 days.

## Methods

### Cell culture and lentiviral NanoLuc® transduction

Human bone marrow-derived hMSCs, passages 3-5, were cultured in Mesenchymal Stem Cell Basal Medium (MSBCM, Lonza, USA). DU145 cell lines were purchased from American Type Culture Collection and cultured in Dulbecco’s Modified Eagle Medium (DMEM, Gibco, USA) supplemented with 10% heat inactivated fetal bovine serum (FBS). To generate Nanoluc-transduced DU145 cells, 5×10^5^ cells were cultured in DMEM/FBS until 80% confluency. Nanoluc luciferase reporter lentiviral vectors were produced by co-transfecting a CMV promotor driven Nanoluc vector (pNL1.1.CMV[Nluc/CMV], cat# N1091, Promega, USA), with VSVG envelope glycoprotein pMD2.G (plasmid #12259) and psPAX2 (plasmid#12260) (Addgene, USA) in 293FT cells. Viral supernatant was concentrated by ultrafiltration and DU145 cells were incubated with the supernatant at (MOI=10) containing polybrene (6 μg/ml). Six hours later, the medium was discarded, and fresh medium was added. To validate Nanoluc transduction, DU145 cells were plated in 6-well plates at densities of 1×10^6^, 1×10^5^, 1×10^4^ and 1×10^3^ cells per well. For in vitro BLI, Nano-Glo® Fluorofurimazine (FFz) powder (Cat#-N4100, Promega, USA) was dissolved in 100% ethanol to yield a 5 mM stock solution and further diluted to 10 μM (1:500) before adding to the wells. The plate was serially imaged with an IVIS Spectrum/CT imaging system (Caliper Life Sciences) every 5 min after substrate addition. 3D tumor spheres were generated using ultra-low attachment 6-well plates (Corning) in DMEM/F12 medium supplemented (Gbco, 11320033) with 10% fetal bovine serum (FBS), 1% penicillin-streptomycin, 1% L-glutamine, 20 ng/mL epidermal growth factor (EGF), and 10 ng/mL basic fibroblast growth factor (bFGF). Cells were seeded at a density of 1×10^4^ cells/well and incubated at 37°C in a 5% CO_2_ humidified atmosphere. Tumor sphere formation was monitored, and spheres reaching a diameter of 200–500 μm were used for all experiments to ensure consistency in size and viability.

### GIONF-labeling of hMSCs

GIONF was prepared as previously described^41^. Prior to labeling, hMSCs were cultured at approximately 70% confluency. For labeling, hMSCs were incubated for 24h with 25 μg Fe/mL GIONF and 3.1 μg/mL PLL (P-1524, Sigma-Aldrich) in 10 ml MSCBM.

### Evaluation of labeling

For PB staining, hMSCs were labeled with GIONF for 24h in 24-well plates on sterilized 15 mm^2^ coverslips. Labeled cells were fixed for 10 minutes with 4% glutaraldehyde,and stained/counterstained with Perl’s reagent/neutral fast red as previously described^42^. Coverslips were washed thrice and embedded on slides with Immuno-Mount. Microscopy was performed using an Apotome 0.2 Zeiss Microscope. For evaluation of cytotoxicity, hMSCs were labeled with 10, 25, 50, or 100 μg Fe/mL GIONF with and without PLL in 96-well plates. An LDH-Cytotoxicity Colorimetric Assay Kit II (MBL International Corporation) was used according to the manufacturer’s protocol. For quantification of cellular iron content, 3×10^3^, 6×10^3^, or 9×10^3^ GIONF-hMSCs were detached and suspended in 50 μL PBS, following which a spectrophotometric Ferrozine assay was performed as previously described^42^.

### Boyden chamber assay

Unlabeled hMSCs and GIONF-hMSCs (1.5×10^5^ cells) were detached, resuspended in 200 μL serum-free DMEM, and seeded into the insert chamber of a 24-well Boyden transwell migration assay (8-μm pore size; Falcon, 353097). The lower chamber contained 500 μL of DU145-conditioned media, RWPE1-conditioned media, or serum-free DMEM as a control. After a 4-hour incubation at 37°C, migrated cells were enzymatically detached from the lower membrane using 0.05% trypsin-EDTA for 5 minutes, followed by three washes in PBS to remove non-adherent cells. Migrated cells were collected, resuspended in 100 μL PBS, and transferred to 400 μL of 0.5% (w/v) agarose in PBS for quantification using MPI. Qualitative migration was assessed via Luna Digital Cell Counter after staining with Acridine Orange / Propidium Iodide (AO/PI) Cell Viability Kit (Logos Biosystem, F23001), following the manufacturer’s recommended protocol.

### MPI/CT studies

For *in vitro* MPI, 1×10^3^, 5×10^3^, 1×10^4^, 5×10^4^, or 1×10^5^ GIONF-hMSCs and 1.25, 2.5, 5.0, and 10.0 μg Fe naked GIONF were suspended first 100 μL PBS, 150 μL 5% agarose gel (Sigma Aldrich) was added and transferred to a 300 μL Eppendorf tube. MPI was performed using a Momentum scanner (Magnetic Insight, Alameda, USA), with image analysis performed using ImageJ software. All animal studies were approved by our Institutional Animal Care and Use Committee. For all procedures, mice were anaesthetized with isoflurane (3–4% induction, 1–2% maintenance). Male immunodeficient rag2^−/−^ mice (6-8 weeks) were obtained from our in-house breeding colony operated under a 12 h light/dark cycle with ad libitum access to food and water. Mice were injected s.c. with 2×10^6^/50 μl Nanoluc-DU145 cells. GIONF-hMSCs (2×10^5^ cells) and 10 μg Fe naked GIONF were injected i.v. or i.t. in DU145-xenografted mice when tumors reached a size of 100-150 mm^3^. Serial 3D MPI and CT was performed 30 min, 24h, 48h, 72h and 7 days post injection. Fiducials (5, 10, and 20% of the injected cell dose) were prepared from the same batches of GIONF-hMSCs and naked GIONFs, serving as MPI quantification references. 2D MPI image analysis was performed using FIJI software and co-registration of CT and 3D MPI using Slicer software. For *ex vivo* MPI, mice were sacrificed on day 7, and the brain, lung, heart, liver, spleen, kidney, and tumor were excised and subjected to 2D MPI/Fiji image analysis.

### PTT and BLI

For *in vitro* PTT, GIONF-hMSCs (1.25×10^4^, 2.5×10^4^, 5×10^4^, 1×10^5^, and 2×10^5^ cells) and naked GIONF (12.5, 25, 50, and 100 μg Fe/50 μL) were suspended in 50 μL PBS and irradiated at 2.2 W/cm^2^ for 150 seconds using an optical fiber connected to a continuous-wave 808 laser source (Changchun New Industries Optoelectronics Tech, China). Temperature monitoring was conducted using a TMI4 Temperature Multichannel Instrument (FISO Technologies Inc., Canada) and infrared images were taken with an FLIR C5 compact thermal camera (Teledyne FLIR, Wilsonville) at 160 × 120 thermal resolution and 5 megapixels. Live/dead cell staining was performed using acridine orange/propidium iodide and analysis with an inverted Zeiss AXIO imager 2 fluorescence microscope. One hundred DU145 3D spheres (sized 300-500 μm) were incubated with 2×10^5^ GIONF-hMSCs or 10 μg of GIONF for 24h in 500 μL of media. Spheres were washed, re-suspended in 50 μL of PBS, and irradiatiated with the same laser at 2.2 W/cm^2^ for 90 seconds. Measurements and live/dead cell staining was performed as described above. For *in vivo* PTT, mice were randomly assigned to 4 experimental groups (n=5 animals each): 1) Control, receiving laser irradiation only (no injection); 2) I.t. injection of 10 μg Fe naked GIONF with irradiation after 30 minutes to determine initial biodistribution; 3) I.t. injection of 10 μg Fe naked GIONF with irradiation after 48h to determine NP retention; and 4) I.t. injection of 2×10^5^ GIONF-hMSCs with irradiation after 48h. All groups were subjected to irradiation at an intensity of 2.2 W/cm^2^ for 7 minutes.

For BLI, 60 μL of Nano-Glo substrate (Cat#-N4100, Promega, Madison WI) was injected i.p. and the mice were imaged with an IVIS Spectrum/CT imaging system before and after performing PTT. Follow-up imaging was conducted at 7, 14, and 30 days. The BLI signal intensity was calculated using Living Imaging software (Caliper Life Sciences).

### Post-mortem tissue analysis

Excised tissues were immersed in 4% paraformaldehyde at 4 °C for 24h, and then transferred to 30% sucrose for another 72h. Liver and lung were cryosectioned at 10 μm slice thickness and stained with H&E and Perl’s reagent.

### Statistics and reproducibility

Data are presented as average values with standard deviation. For statistical analysis involving multiple groups, a one-way ANOVA was used, followed by Dunnett’s test for specific comparisons. Significance is indicated as *P<0.05, **P<0.01, ***P<0.001 and ****P<0.0001. For two-group comparisons, a two-tailed Student’s t-test was employed, with the same significance thresholds. The number of independent experimental repetitions is given in the corresponding figure legends. All statistical analysis was carried out using GraphPad Prism version 9.0. All researchers conducting the experiments and processing MPI, BLI and PTT data were blinded to the experimental group assignments.

### Reporting summary

Further information on research design is available in the Nature Portfolio Reporting Summary linked to this article.

## Data availability

The data that support the findings of this study are available within the article and Supplementary Information and Supplementary Data. The raw imaging data are available from the corresponding author upon reasonable request.

### Acknowledgements

This work was supported by research grants from the NIH (P41 EB024495, R01 EB030376, R01 CA257557, UH2/UH3 EB028904 and S10 OD026740 to J.W.M.B), the Maryland Stem Cell Research Fund (MSCRFD-5416 to J.W.M.B.), and the French National Agency of Research (ANR-23-CE18-0016-01 to S.R.).

## Author contributions

B.G., A. S-Z. and J.W.M.B. conceived and designed the research. B.G. and A.R. performed all *in vitro* and *in vivo* animal experiments and data analysis. G.L, R.B. and S.R. performed synthesis of the GIONF nanoparticles. S.K assisted with lentiviral transduction. B.G. and A.R. wrote the manuscript draft. J.W.M.B. supervised the whole project. All authors have read and revised the manuscript.

## Additional information

## Supplementary information

The online version contains supplementary material available at https://doi.org/….

## Correspondence and requests for materials

should be addressed to Jeff W.M. Bulte.

## References

1. Chen, Q. et al. Recent advances in different modal imaging-guided photothermal therapy. Biomaterials 106, 144–166 (2016).

2. Riley, R.S. & Day, E.S. Gold nanoparticle-mediated photothermal therapy: applications and opportunities for multimodal cancer treatment. Wiley Interdiscip Rev Nanomed Nanobiotechnol 9 (2017).

3. Arami, H. et al. Remotely controlled near-infrared-triggered photothermal treatment of brain tumours in freely behaving mice using gold nanostar s. Nature nanotechnology 17, 1015–1022 (2022).

4. Wang, L. et al. Exploring and Analyzing the Systemic Delivery Barriers for Nanoparticles. Adv Funct Mater 34 (2024).

5. Salimi, M., Kuddannaya, S. & Bulte, J.W.M. Pharmacokinetic Profiling of Unlabeled Magnetic Nanoparticles Using Magnetic Particle Imaging as a Novel Cold Tracer Assay. Nano Lett (2024).

6. Schweizer, M.T. et al. A Phase I Study to Assess the Safety and Cancer-Homing Ability of Allogeneic Bone Marrow-Derived Mesenchymal Stem Cells in Men with Localized Prostate Cancer. Stem Cells Transl Med 8, 441–449 (2019).

7. Stuckey, D.W. & Shah, K. Stem cell-based therapies for cancer treatment: separating hope from hype. Nature Reviews Cancer 14, 683–691 (2014).

8. Rosu, A., Ghaemi, B., Bulte, J.W.M. & Shakeri-Zadeh, A. Tumor-tropic Trojan horses: Using mesenchymal stem cells as cellular nanotheranostics. Theranostics 14, 571–591 (2024).

9. Xu, L. et al. Quantitative Comparison of Gold Nanoparticle Delivery via the Enhanced Permeation and Retention (EPR) Effect and Mesenchymal Stem Cell (MSC)-Based Targeting. ACS Nano 17, 2039–2052 (2023).

10. Lai, Y.H. et al. Stem cell-nanomedicine system as a theranostic bio-gadolinium agent for targeted neutron capture cancer therapy. Nat Commun 14, 285 (2023).

11. Huang, X. et al. Design considerations of iron-based nanoclusters for noninvasive tracking of mesenchymal stem cell homing. ACS Nano 8, 4403–4414 (2014).

12. Shakeri-Zadeh, A. & Bulte, J.W.M. Imaging-guided precision hyperthermia with magnetic nanoparticles. Nat. Rev. Bioeng. 2, 10.1038/s44222-44024-00257-44223 (2024).

13. Bulte, J.W.M. Superparamagnetic iron oxides as MPI tracers: A primer and review of early applications. Adv Drug Deliv Rev 138, 293–301 (2019).

14. Rastinehad, A.R. et al. Gold nanoshell-localized photothermal ablation of prostate tumors in a clinical pilot device study. Proc Natl Acad Sci U S A 116, 18590–18596 (2019).

15. Luo, M. et al. Mesenchymal stem cells transporting black phosphorus-based biocompatible nanospheres: Active trojan horse for enhanced photothermal cancer therapy. Chemical Engineering Journal 385, 123942 (2020).

16. Bulte, J.W. et al. Quantitative “Hot Spot” Imaging of Transplanted Stem Cells using Superparamagnetic Tracers and Magnetic Particle Imaging (MPI). Tomography 1, 91–97 (2015).

17. Cromer Berman, S.M. et al. Cell motility of neural stem cells is reduced after SPIO-labeling, which is mitigated after exocytosis. Magn Reson Med 69, 255–262 (2013).

18. Bulte, J.W.M., Wang, C. & Shakeri-Zadeh, A. In Vivo Cellular Magnetic Imaging: Labeled versus Unlabeled Cells. Advanced Functional Materials 32, 2207626 (2022).

19. Brachi, G. et al. Intratumoral injection of hydrogel-embedded nanoparticles enhances retention in glioblastoma. Nanoscale 12, 23838–23850 (2020).

20. Yu, M., Zhang, C., Tang, Z., Tang, X. & Xu, H. Intratumoral injection of gels containing losartan microspheres and (PLG-g-mPEG)-cisplatin nanoparticles improves drug penetration, retention and anti-tumor activity. Cancer Letters 442, 396–408 (2019).

21. Huang, L. et al. Intelligent photosensitive mesenchymal stem cells and cell-derived microvesicles for photothermal therapy of prostate cancer. Nanotheranostics 3, 41 (2019).

22. Xu, C. et al. A Light-Triggered Mesenchymal Stem Cell Delivery System for Photoacoustic Imaging and Chemo-Photothermal Therapy of Triple Negative Breast Cancer. Adv Sci (Weinh) 5, 1800382 (2018).

23. Xiao, J. et al. Tumor-Tropic Adipose-Derived Mesenchymal Stromal Cell Mediated Bi(2) Se(3) Nano-Radiosensitizers Delivery for Targeted Radiotherapy of Non-Small Cell Lung Cancer. Adv Healthc Mater 11, e2200143 (2022).

24. Yamamoto, A. et al. Metastasis from the tumor interior and necrotic core formation are regulated by breast cancer-derived angiopoietin-like 7. Proc Natl Acad Sci U S A 120, e2214888120 (2023).

25. Shi, J., Kantoff, P.W., Wooster, R. & Farokhzad, O.C. Cancer nanomedicine: progress, challenges and opportunities. Nature Reviews Cancer 17, 20–37 (2017).

26. Mooney, R. et al. Neural stem cell-mediated intratumoral delivery of gold nanorods improves photothermal therapy. ACS Nano 8, 12450–12460 (2014).

27. Pampaloni, F., Reynaud, E.G. & Stelzer, E.H.K. The third dimension bridges the gap between cell culture and live tissue. Nature Reviews Molecular Cell Biology 8, 839–845 (2007).

28. Weiswald, L.-B., Bellet, D. & Dangles-Marie, V. Spherical Cancer Models in Tumor Biology. Neoplasia 17, 1–15 (2015).

29. Nicolás-Boluda, A. et al. Photothermal Depletion of Cancer-Associated Fibroblasts Normalizes Tumor Stiffness in Desmoplastic Cholangiocarcinoma. ACS Nano 14, 5738–5753 (2020).

30. Chen, A.L. et al. Quantifying spectral changes experienced by plasmonic nanoparticles in a cellular environment to inform biomedical nanoparticle design. Nanoscale Research Letters 9, 454 (2014).

31. Lartigue, L. et al. Cooperative Organization in Iron Oxide Multi-Core Nanoparticles Potentiates Their Efficiency as Heating Mediators and MRI Contrast Agents. ACS Nano 6, 10935–10949 (2012).

32. Liu, Y. et al. Glutathione-Responsive Self-Assembled Magnetic Gold Nanowreath for Enhanced Tumor Imaging and Imaging-Guided Photothermal Therapy. ACS Nano 12, 8129–8137 (2018).

33. Li, C. et al. Gold-Coated Fe(3)O(4) Nanoroses with Five Unique Functions for Cancer Cell Targeting, Imaging and Therapy. Adv Funct Mater 24, 1772–1780 (2014).

34. Xiao, J. et al. Near Infrared-Absorbing Nanoparticle-Mediated Endovascular Photothermal Precision Embolization of Tumor Feeding Vessels for Starvation Treatment. ACS Biomaterials Science & Engineering 9, 3660–3669 (2023).

35. Mooney, R. et al. Neural stem cell-mediated intratumoral delivery of gold nanorods improves photothermal therapy. ACS nano 8, 12450–12460 (2014).

36. Li, X., Lovell, J.F., Yoon, J. & Chen, X. Clinical development and potential of photothermal and photodynamic therapies for cancer. Nat Rev Clin Oncol 17, 657–674 (2020).

37. Kang, S. et al. Mesenchymal Stem Cells Aggregate and Deliver Gold Nanoparticles to Tumors for Photothermal Therapy. ACS Nano 9, 9678–9690 (2015).

38. Kim, J.-W., Galanzha, E.I., Shashkov, E.V., Moon, H.-M. & Zharov, V.P. Golden carbon nanotubes as multimodal photoacoustic and photothermal high-contrast molecular agents. Nature nanotechnology 4, 688–694 (2009).

39. Bourrinet, P. et al. Preclinical Safety and Pharmacokinetic Profile of Ferumoxtran-10, an Ultrasmall Superparamagnetic Iron Oxide Magnetic Resonance Contrast Agent. Investigative Radiology 41 (2006).

40. Balasubramanian, S.K. et al. Biodistribution of gold nanoparticles and gene expression changes in the liver and spleen after intravenous administration in rats. Biomaterials 31, 2034–2042 (2010).

41. Hugounenq, P. et al. Iron Oxide Monocrystalline Nanoflowers for Highly Efficient Magnetic Hyperthermia. The Journal of Physical Chemistry C 116, 15702–15712 (2012).

42. Bulte, J.W., Arbab, A.S., Douglas, T. & Frank, J.A. Preparation of magnetically labeled cells for cell tracking by magnetic resonance imaging. Methods Enzymol 386, 275–299 (2004).

