## Supplemental Figures for "MPI-Guided Photothermal Therapy of Prostate Cancer using Stem Cell Delivery of Magnetotheranostic Nanoflowers"

*[1]The Russell H. Morgan Department of Radiology and Radiological Science, The Johns Hopkins University School of Medicine, Baltimore, MD 21205, USA; [2]Cellular Imaging Section and Vascular Biology Program, Institute for Cell Engineering, The Johns Hopkins University School of Medicine, Baltimore, MDd 21205, USA; [3]Université de Franche-Comté, CNRS, Chrono-environnement, F-25000, Besançon, France; [4]Department of Biomedical Engineering, Johns Hopkins University, Baltimore, MD21205, USA; [5]Department of Chemical & Biomolecur Engineering, Johns Hopkins University Whiting School of Engineering, Baltimore, MD 21218; [6]Department of Oncology, Johns Hopkins University, Baltimore, MD 21287, USA; [7]F. M. Kirby Research Center for Functional Brain Imaging, Kennedy Krieger Inc., Baltimore, MD 21205, USA.*

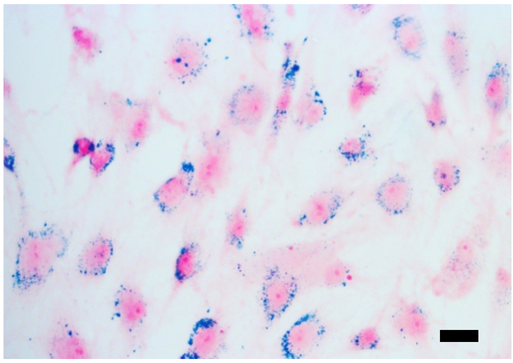

**Fig. S1.** Prussian blue staining of hMSCs incubated for 24 hours with 25 μg Fe/mL GIONF without PLL demonstrates intracellular peri-nuclear labeling. Scale bar=10 μm.

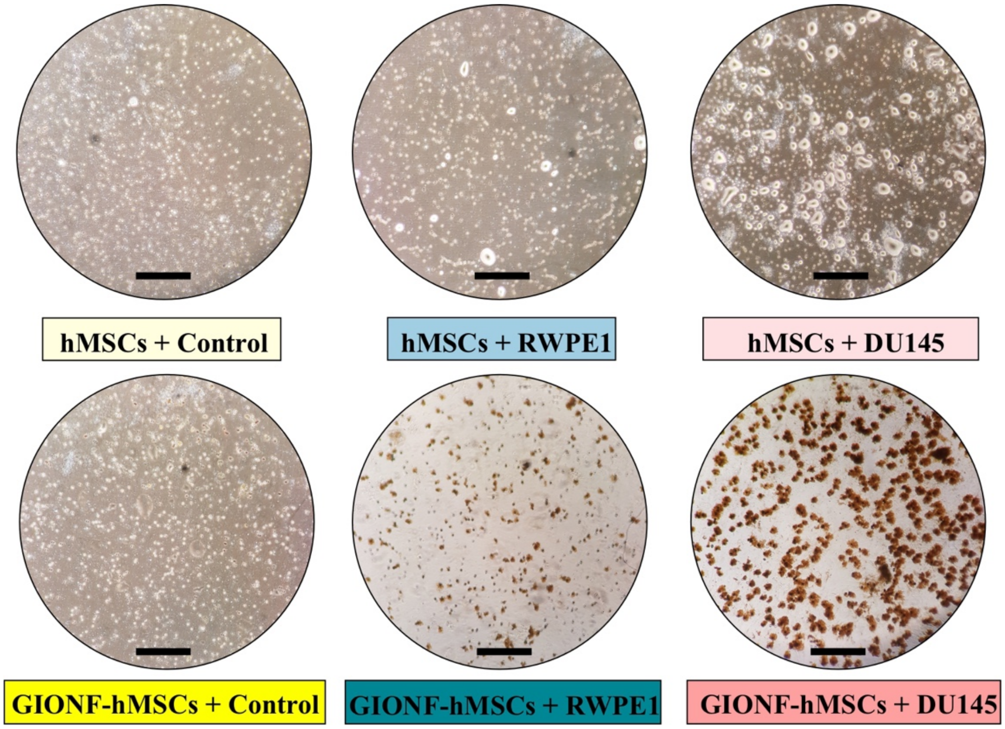


**Fig. S2.** Bright-field microscopy of hMSC and GIONF-hMSC tumor tropism for DU145 cells compared to control medium (no cells) and normal prostate (RWPE1) cells. Scale bar=50 μm.

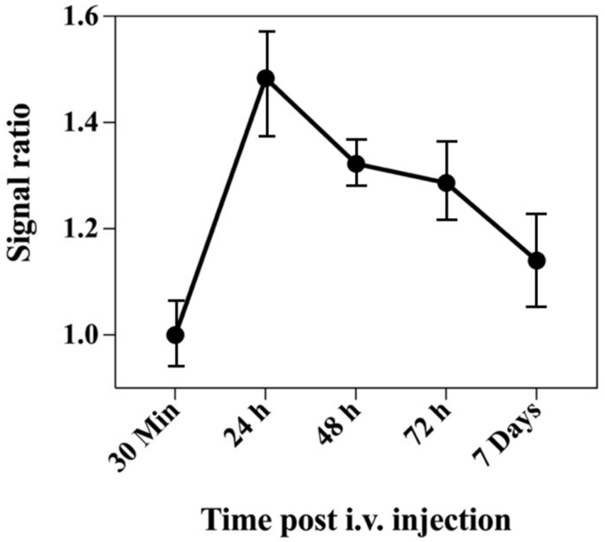


**Fig. S3.** MPI liver signal ratios of the MPI signal intensity at each time point divided by the initial MPI signal at 30 min post i.v. injection of GIONF-hMSCs. (n=3 animals).

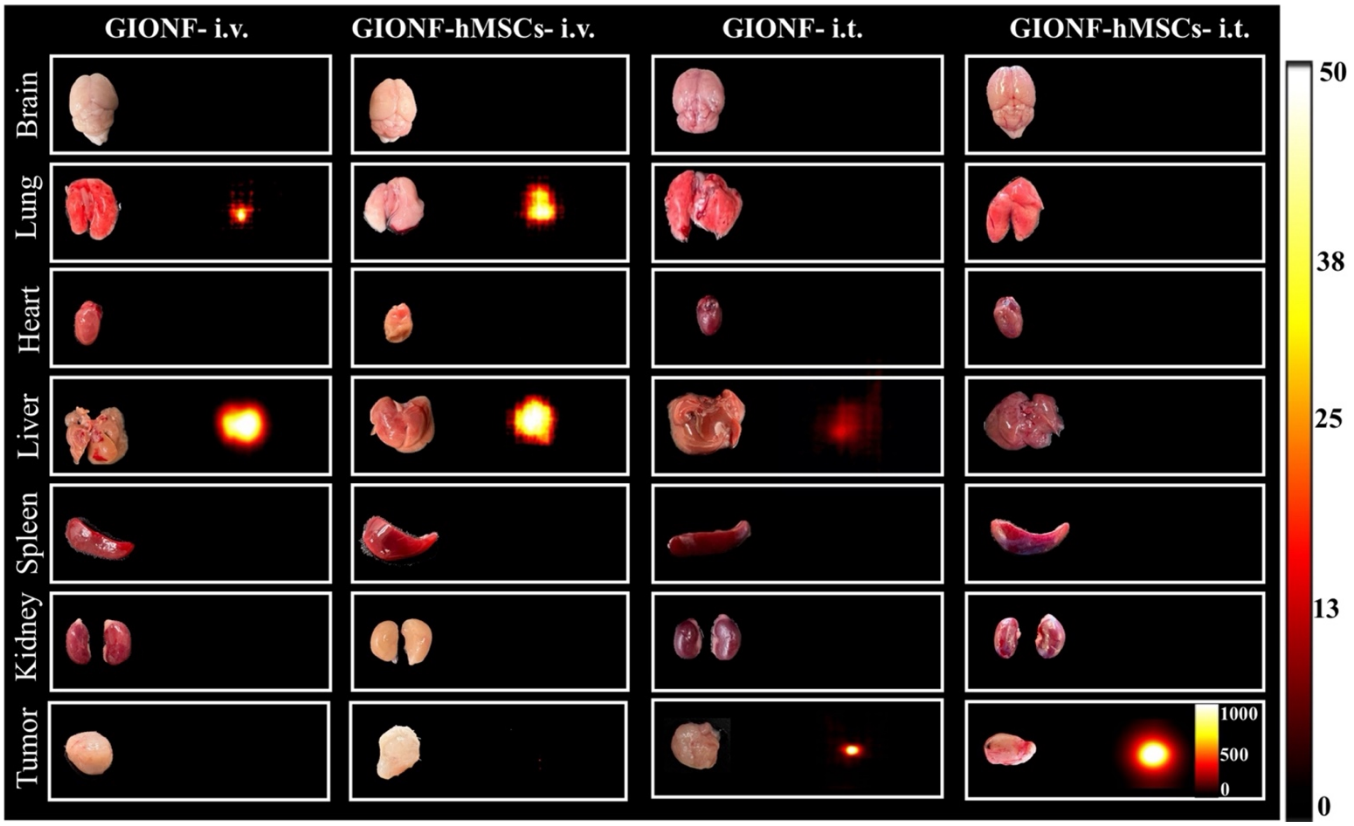


**Fig. S4.** *Ex vivo* 2D MPI of the tumor and other organs 7 days post i.v. or i.t. injection of naked GIONF or GIONF-hMSCs.

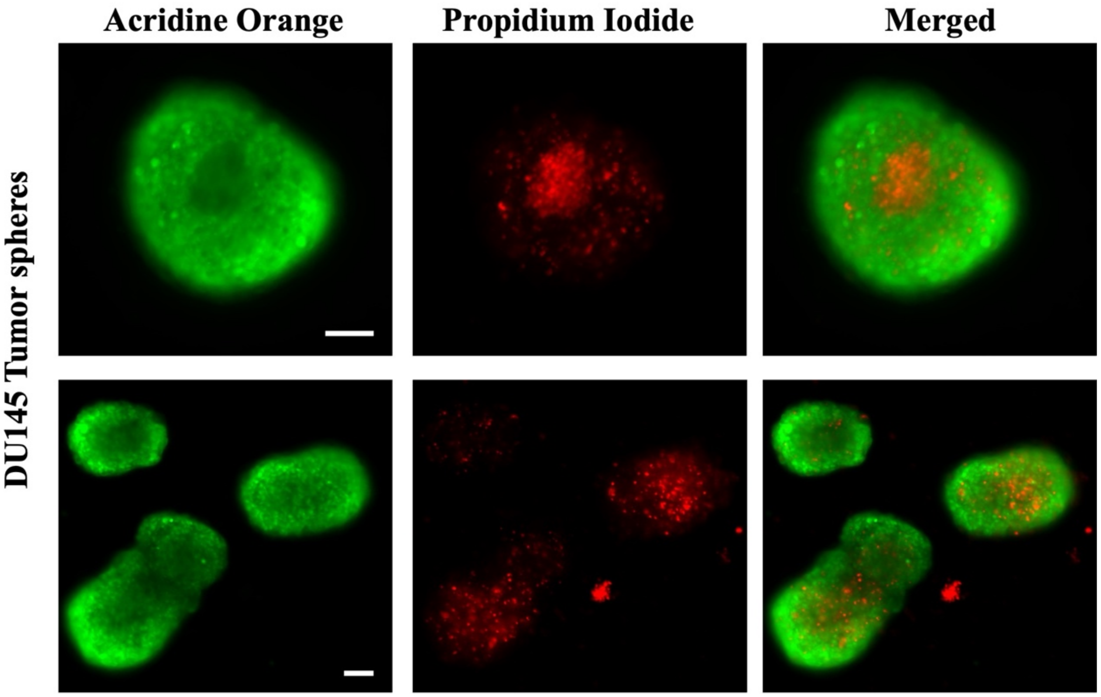


**Fig. S5.** Fluorescent staining of tumor spheres with using Acridine Orange and Propidium Iodide (PI). The image reveals the characteristic zonation of large spheroids, including an outer proliferative layer, and a distinct necrotic core as visualized by PI uptake. Scale bar=100 μm.

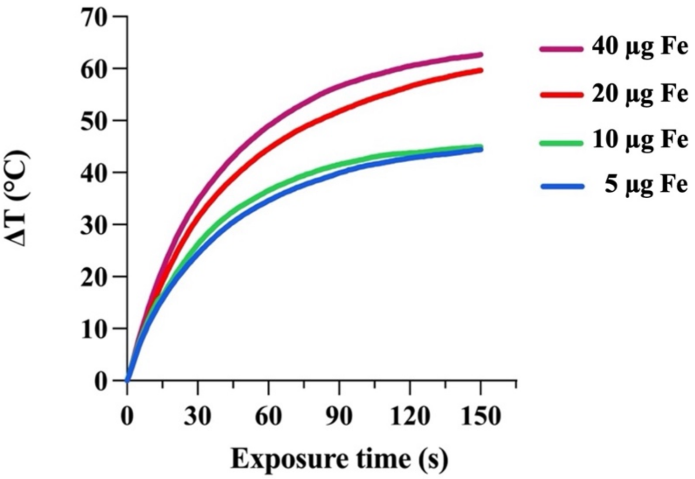


**Fig. S6.** Thermometry of naked GIONF solutions containing different amounts of Fe exposed to an 808 nm NIR laser (2.2 W/cm$^2$, 150 s).
